# Contribution of Wide Dynamic Range neuronal activation to somatosensory evoked potentials supports use as biomarker of spinal nociceptive processing

**DOI:** 10.64898/2026.07.16.738872

**Authors:** Kenneth Steel, Tony Blockeel, Elise Ajay, Jeff Krajewski, Keith Phillips, James P Dunham, Anthony E Pickering

**Affiliations:** University of Bristol; University of Bristol School of Physiology Pharmacology and Neuroscience; Eli Lilly; Grunenthal

## Abstract

Identifying objective translational biomarkers of spinal nociceptive processing is important to accelerate analgesic development. The primary negative component (N1) of spinal somatosensory evoked potentials (SEPs) has been proposed as such a biomarker. However, the cellular substrates of the N1 potential (evoked by innocuous electrical stimulation) and their relevance to nociceptive processing have not been directly demonstrated. Here, we employed a 64-channel multielectrode recording approach in the dorsal horn of anaesthetised Wistar rats to functionally characterise the neuronal populations activated during the generation of spinal SEPs and determine how their activity is modulated by tapentadol. Single units were classified based on their responses to mechanical stimulation of the hindpaw, and their electrically evoked responses to sciatic nerve stimulation. Of 59 well-isolated units, 47 (80%) were classified as wide dynamic range (WDR) neurons and 12 (20%) as low-threshold mechanoreceptive (LTMR) neurons, spatially distributed across spinal laminae III-V. Tapentadol (10 mg/kg, intraperitoneal (i.p)) selectively attenuated the mechanically- and electrically-evoked activity of WDR neurons without affecting LTMR responses. This WDR-inhibition was largely reversed by naloxone (0.25 mg/kg, i.p) but not by atipamezole (1 mg/kg, i.p), identifying a predominant opioid receptor-mediated mechanism of inhibition in the naïve state. The magnitude of WDR inhibition by tapentadol correlated with the degree of reduction of the N1 amplitude. These findings establish activity in WDR neurons as a core component of the N1 potential, supporting the use of spinal SEPs as a translational biomarker of analgesic target engagement within the dorsal horn.

**Summary:** Multielectrode recordings identify inhibition of WDR neurons as the mechanism by which tapentadol modulates spinal SEPs, supporting use as a biomarker of spinal nociceptive processing.

## Introduction

Objective translatable biomarkers of spinal nociceptive processing could help to accelerate the development of analgesics which target specific mechanisms in chronic pain conditions [27]. We have previously shown in a reverse translational study that the amplitude of the primary negative component (N1) of spinal somatosensory evoked potentials (SEPs) is differentially modulated by analgesics in a manner consistent with their distinct mechanisms of action. This provided proof-of-concept evidence that SEPs could serve as a biomarker of analgesic target engagement within the spinal cord [37]. The generation of the N1 component (termed N13 in humans) has been attributed to post-synaptic activity driven by Aβ primary afferent inputs. Since most Aβ fibres are activated by innocuous stimuli such as light brushing, their post-synaptic targets in the dorsal horn were not considered relevant to nociceptive processing [5,9,12].

An emerging hypothesis challenging this view proposes that the N1 component could also reflect the post-synaptic activation of wide dynamic range (WDR) neurons [23,30]. WDR neurons integrate convergent input from many classes of primary afferents — encoding stimulus intensity across the full innocuous-to-noxious range. WDR neurons are strongly implicated in the process of central sensitisation following peripheral nerve injury, where their heightened responsiveness has been reported to be a feature of the development of neuropathic pain [38,44]. Several human pain studies investigating the sensitivity of the N13 component to interventions using experimental models support this proposition. In a conditioned pain modulation paradigm, cold pressor stimulation significantly attenuated N13 amplitude [30], an effect attributed to the engagement of endogenous pain modulatory systems acting upon WDR neurons. In another study, capsaicin-induced an increase of the N13 amplitude which was prevented by pre-treatment with pregabalin [25], consistent with suppression of sensitisation at the spinal cord [4,16,26]. These findings support the idea that changes in the N13 component could reflect changes in the excitability of WDR neurons. However, the non-invasive recording techniques used in these studies precludes the direct attribution of N13 amplitude changes to specific neuronal populations.

It remains unknown which populations of spinal neurons are the cellular targets of analgesics acting to suppress the amplitude of the innocuously evoked N1 component, and whether these neurons contribute to the encoding of noxious stimuli. To address this directly, we employed a multielectrode recording approach in the rat dorsal horn to characterise functional neuronal populations activated during spinal SEP generation. We then determined how their responses to electrical and mechanical stimulation are modulated by tapentadol. We identified low-threshold mechanoreceptive (LTMR) and WDR neurons as contributors to the N1 component. Tapentadol selectively attenuated the mechanically- and electrically-evoked activity of WDR neurons without affecting LTMR responses via an opioidergic action. The degree of WDR inhibition correlated with the change in amplitude of the N1 component, establishing WDR neurons as a key neuronal component of the N1 potential and its modulation by tapentadol.

## Methods

### Animals

All experiments were conducted in accordance with the UK Animals (Scientific Procedures) Act 1986 and approved by the University of Bristol Animal Welfare and Ethical Review Board. Male Wistar rats (n=12, Envigo, UK) weighing between 225g-275g were housed in cages under a 12-hour light/dark cycle with *ad libitum* access to food and water.

### Experimental preparation

#### Drugs

Each rat received intraperitoneal injections of tapentadol (10mg/kg, Precise Chemipharma, India) and either naloxone (0.25mg/Kg, Tocris, Bristol, UK) or atipamezole (1mg/kg, Tocris, Bristol, UK) to a volume of 10ml/kg in 0.9% saline.

#### Surgery

Anaesthesia was induced with isoflurane until loss of the paw withdrawal reflex (5% isoflurane at 1.0L/min). The rat was transferred to a preparation area where a deep plane of anaesthesia was maintained via a nose cone (2% isoflurane at 0.5L/min). The skin over the thoracolumbar spine and the bicep femoris muscle was shaved and sterilised with chlorohexidine wipes. A custom made canula was inserted into the lower right quadrant of the peritoneal cavity for drug delivery. The rat was then positioned in the ear bars of a stereotaxic frame (David Kopf Instruments, California, USA), on top of a heat mat with rectal temperature monitoring to maintain a core body temperature of 37°C (Harvard Apparatus, Massachusetts, USA).

Following a skin incision above the T13-L1 spinal vertebrae and blunt dissection of the connective tissue and paraspinal muscles, rongeurs were used to expose the dorsal surface of the L4-L5 spinal cord segment. The spine was stabilised for electrophysiology in vertebral clamps attached to the spinous processes above and below the recording site. The tip of a 25G needle was used to incise the dura and pia, and a drop of mineral oil was applied to prevent the tissue from drying.

To access the sciatic nerve a 2cm skin incision was made over the left bicep femoris and careful blunt dissection along the muscle fibres exposed the sciatic nerve proximal to its trifurcation. The skin edges were retracted by suturing to an O-ring, creating a bath for mineral oil (Figure 1A-B). To elicit spinal SEPs, electrical stimulation was delivered directly to the sciatic nerve via a custom-made bipolar Ag-AgCl hook electrodes. To elicit spinal SEPs during the 4Hz electrical stimulation, the stimulus intensity was set to 1.5x the motor threshold (as is convention in human studies) for hindpaw twitch. This was determined in each animal by delivering a 0.2ms electrical pulse every 10s, starting at 0.05mA and incrementally increasing by 0.05mA until the threshold was identified. The intensity, duration and frequency of the electrical stimuli were controlled from Spike2 software controlling a Micro1401 MKII A-D converter (both Cambridge Electronic Design, Cambridge UK) and generated using a DS4 isolated current stimulator (Digitimer Ltd, Welwyn Garden City, UK).

**Figure 1.**
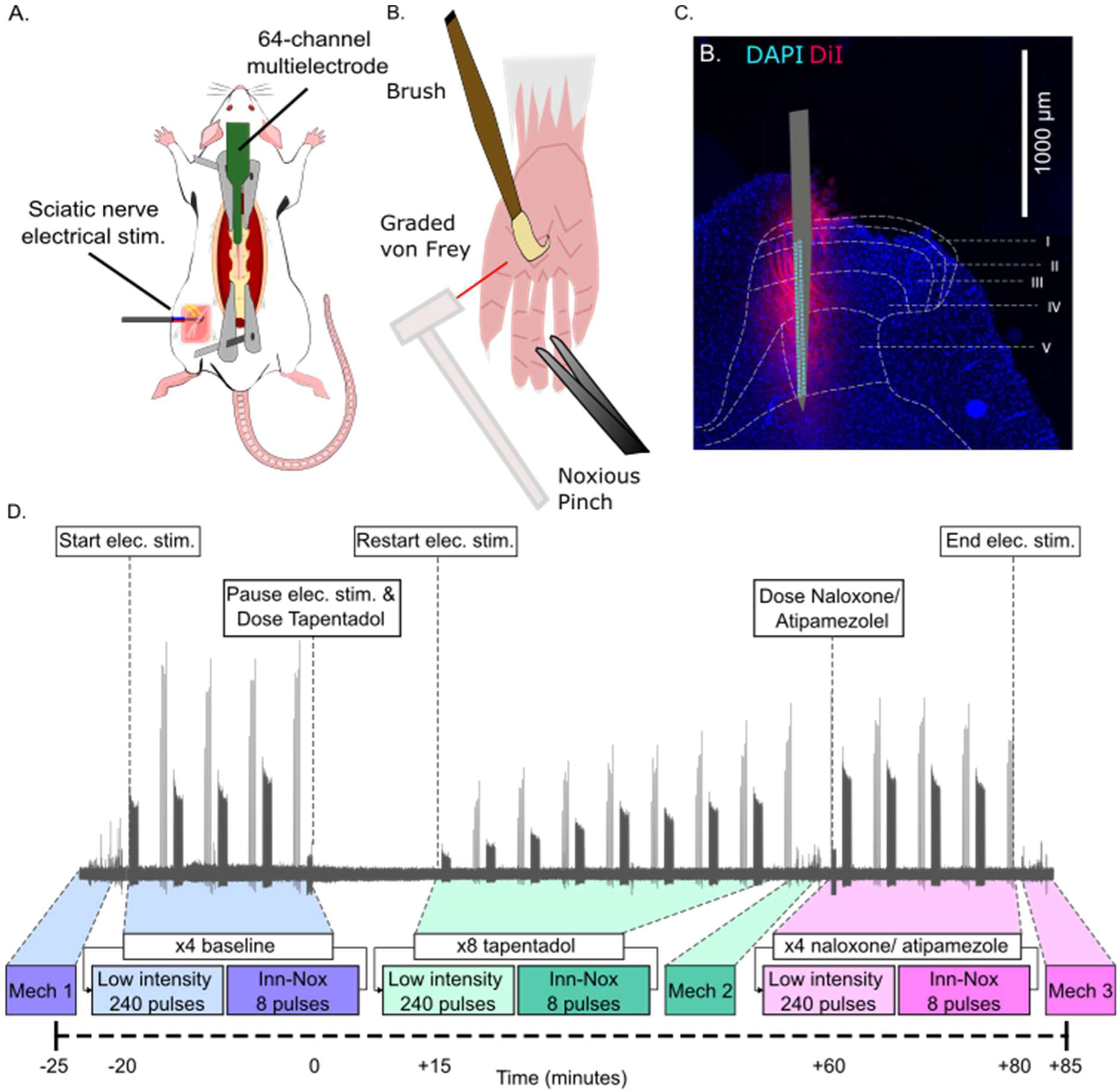
Overview of experimental approach. A. Following exposure of the L4/5 spinal cord and sciatic nerve, a multielectrode silicon probe was inserted to the spinal dorsal horn to record spinal neuronal activity evoked by electrical stimulation of the nerve. B. Confirmation of correct probe position was obtained by observing graded neuronal responses to mechanical stimulation of the receptive field on the hind paw.C. Recording track within spinal cord marked with DiI applied to the probe prior to insertion. D. Recording protocol and exemplar local field potentials recorded from a single contact of the multielectrode silicon probe. Spinal SEPs and mechanically evoked activity were recorded at baseline (blue), following 10mg/kg tapentadol (green), and after an injection of either 0.25mg/kg naloxone or 1mg/kg atipamezole (pink). The electrical stimulation protocol was repeated 4 times at baseline, 8 times after tapentadol and a further 4 times after the administration of naloxone or atipamezole.

#### Spinal recordings

Neural activity was recorded with a 64-channel multicontact silicon probe (ASSY-77 H5, Cambridge Neurotech, Cambridge, UK) connected to 2x 32 channel amplifiers (RHD2132, Intan, California, USA) with a 64-channel adaptor (Neuronexus, Michigan, USA). Data were sampled at 30kHz, band passed filtered and visualised both at 1-300Hz (field potentials) and 600-6000Hz using an acquisition system and GUI (Open Ephys, Atlanta, USA). The multicontact probe was inserted into the spinal dorsal horn (10µm/s) using a single axis motorised manipulator (Scientifica, Uckfield, UK) mounted on the stereotaxic manipulator and frame. It was lowered until spontaneous and mechanically-evoked action potentials (via light brush of the hind paw) were observed along the length of the probe. The depth of the probe was then adjusted to position the maximal amplitude of the electrically evoked N1 component in the central channels of the probe. In some experiments the recording site was marked with DiI lipophilic tracer (Thermo Fisher Scientific, Massachusetts, USA), by submerging the probe in the solution prior to insertion (Figure 1C).

#### Recording protocol

Once the setup for recording spinal SEPs and multiunit neuronal activity was established, a 110-minute recording protocol was initiated - only one recording session was undertaken per animal (Figure 1D). The protocol was designed to generate spinal SEPs with:

1. innocuous electrical stimulation at a 4Hz frequency (240 pulses per train) (Low intensity), and
2. with step-incremented electrical stimuli increasing from the innocuous to the noxious range (Inn-Nox) [37].

Additionally, the protocol included mechanical stimulation of the receptive field which allowed the functional classification of single units based upon their response properties. Mechanical stimulation sets consisted of 3 repeats of light brush, three repeats of von Frey filament application at each weight (0.6, 1, 2, 4, 8, 15 and 26g) and a single pinch applied with blunt forceps. Each of these stimulus sets were performed at baseline, following an i.p injection of 10mg/kg tapentadol, and after an injection of either naloxone (0.25mg/kg, n=5 rats) or atipamezole (1mg/kg, n=7 rats).

#### Histology

In experiments where the recording site was marked with a fluorescent marker (DiI), the animal was perfused with phosphate buffered saline (PBS) and the spinal cord obtained from the vertebral column via hydraulic extrusion (Richner et al., 2017). Spinal tissue was immersion fixed in a solution of 4% paraformaldehyde for 24 hours before being transferred into a 30% sucrose solution for cryoprotection and stored at 4°C. The lumbar spinal cord was position vertically in Cryomatrix (ThermoFisher, UK) on the stage of a freezing Peltier, before being cut with a microtome into 50µm sections. Spinal sections were mounted in series onto microscope slides (Epredia Superfrost, UK) and left to dry for 24 hours. To identify the cytoarchitecture of the spinal dorsal horn, 200µl of a 250nM DAPI (4’,6-diamidino-2-phenylindole) solution was applied to the dried spinal sections for 5 minutes before rinsing to fluorescently tag cell nuclei. The sections were cover slipped in FluoroSave (Sigma-Aldrich, Gillingham, UK).

All sections were viewed under an epifluorescence microscope (Leica DM 4000B, Leica Camera, Wetzlar Germany) and imaged at a 5× magnification. To identify the position of the probe, sections were illuminated with green light (Excitation 545nm, dichroic mirror 562nm, emission 605nm) to reveal the DiI staining. An ultraviolet filter set (excitation 365nm, dichroic mirror 405nm, emission 445nm) was used to visualise the DAPI staining of nuclei. All images were visualised in Fiji [35] and merged to a create compound picture of DAPI and DiI fluorescence. Probe position reconstruction was performed using a custom-written plugin (Wolfson Bioimaging Facility, UK) and the boundaries of Rexed laminae were fitted onto the image according to the L5 segment in the Rat Spinal Cord Atlas [42].

### Data analysis

#### Automated clustering of recorded units

Data was pre-processed by bandpass filtering (300-6000Hz) to show the individual spikes (Figure 2A). The clustering of multielectrode data was performed using the Kilosort4 Python package [29] chosen because of its ability to resolve spike waveforms from nonstationary recordings (drift) and background noise. After pre-processing, including electrical stimulus artefact removal whereby data around each event marker (2ms before to 5ms after) was set to the mean voltage from the entire recording, and all 64 channels of data were merged into a single binary file for Kilosort4, using a custom MATLAB script. Clustering was performed using the standard default settings, except for: the number of channels (n_chan_bin = 64 to match the number of electrode contacts on the probe), the vertical and horizontal spacing of electrode contacts (dmin = 22.5µm & dminx = 12.5µm to match the probe) and the number of template groupings (x_centers = 4 as recommended when using multielectrode probes arranged in a 2D-grid).

**Figure 2.**
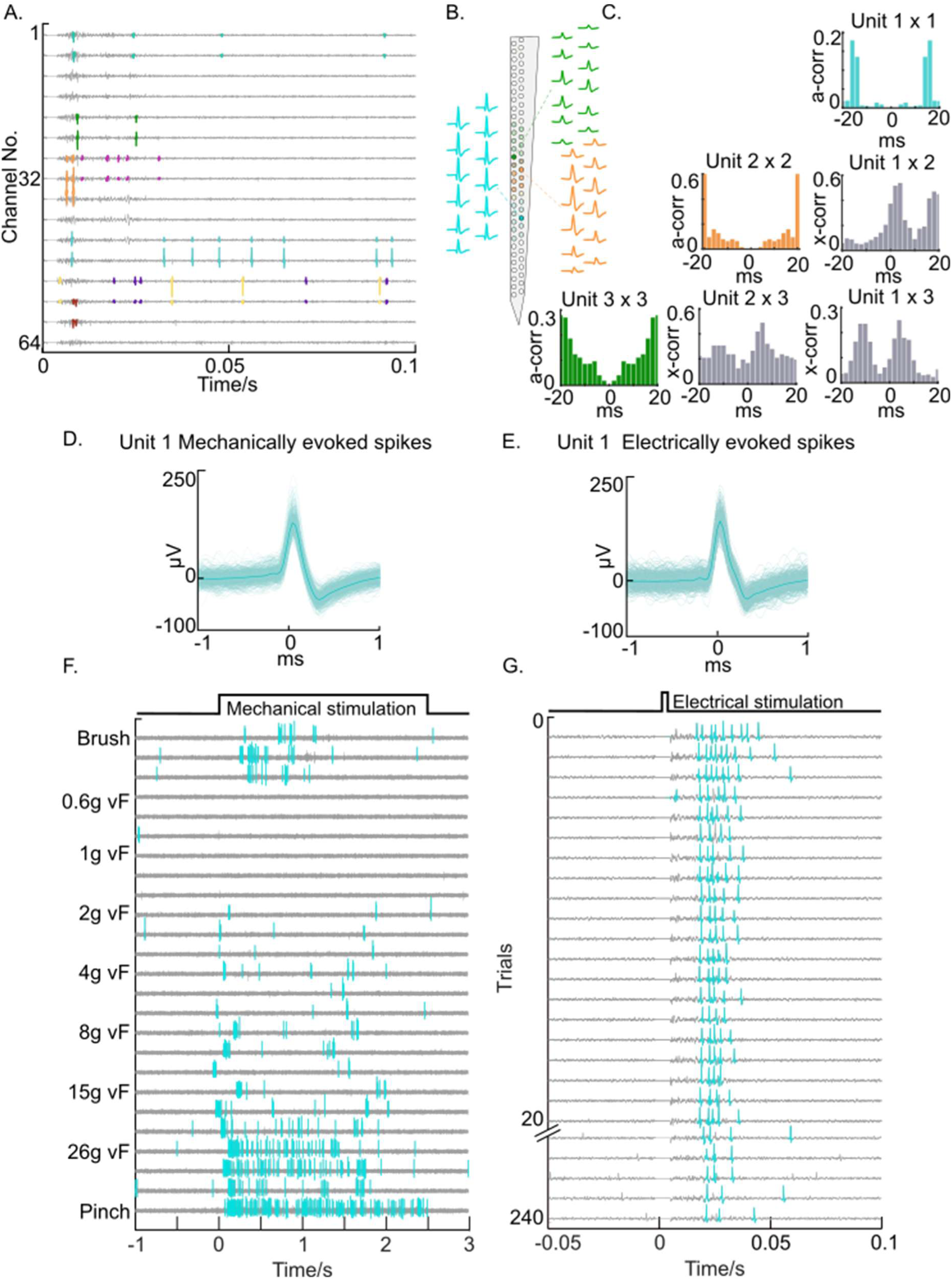
Automated and manual clustering of multielectrode data. A. Example of multielectrode recordings in the spinal dorsal horn following electrical stimulation of sciatic nerve at time 0. Spikes from simultaneously recorded neurons are highlighted along the length of the probe and colour coded into single clusters. B. Putative single units were visualised to ensure their action potentials had a consistent waveform morphology that was visible across multiple electrode contacts. C. Auto- and cross-correlograms the activity of individual neurons (evidenced by high correlation in the cross-correlograms, reflecting synchronised spiking caused by confirm the spiking activity of each cluster was well isolated from multiunit activity (evidenced by ±2ms refractory period in the auto-correlograms) and that they represented electrical stimulation). D. For each unit, the mechanically and E. electrically evoked spikes were superimposed and averaged, to ensure the clusters only contained action potentials with a consistent spike waveform across stimulus modalities. F. Single units were classified based on their responses to mechanical stimulation, before G. their responses to electrical stimulation were assessed.

#### Manual cluster curation

The putative clusters identified automatically by Kilosort4 were manually curated in Phy2 to ensure they only contained spiking activity from a single neuron. This approach closely followed the Phy2 Manual Clustering Guide (https://github.com/cortex-lab/phy) and involved the merging/splitting of clusters as necessary. Clusters were classified as “*good*” if:

1. They had an action potential waveform that was recorded across multiple electrode contacts, with a large amplitude and a consistent waveform (Figure 2A-B).
2. They displayed a clear refractory valley (of width ≥ ±2ms) around time 0 in the auto-correlogram view and showed increased correlation with other clusters on nearby channels in the cross-correlogram view (Figure 2C).
3. The activity of the neuron could be tracked throughout the entire duration of the recording.

Clusters that were missing spikes from a section of the recording were either merged with clusters that contained the missing spikes (evidenced by an identical waveform morphology and a lack of refractory period violations in the cross-correlogram view), or they were excluded from further analysis. Equally, clusters that appeared to contain action potentials from multiple neurons were split into individual neurons where possible, to be either merged with other well-isolated clusters, or classified as “*multiunit activity*” and excluded from further analysis.

#### Functional classification and single unit analysis

Visualisation and analysis of single units was performed in MATLAB 2022b, with custom scripts available on <u>GitHub</u> and adapted from [34]. The cluster spike times and event times of the electrical and mechanical stimulation were imported into MATLAB along with the .continuous file containing the raw data from each electrode contact. Auto- and cross-correlograms of the clusters were generated again in MATLAB using the ‘xcorr’ function, to calculate the auto- and cross-correlation of spike times in 2ms bins across the entire length of the recording. The spike times and filtered data were used to generate action potential waveforms (covering ±1ms of each spike time) of single unit responses to electrical and mechanical stimulation, to confirm clusters contained spikes with consistent waveform morphology across different stimulation modalities (Figure 2F-G). The electrode contacts which recorded the peak action potential waveform was used to estimate the depth and hence putative spinal dorsal horn lamina where each unit was positioned.

The identified isolated units were classified into functional populations based on their response characteristics to mechanical stimulation at baseline. Raster plots were generated to display the mechanically evoked activity of each neuron by plotting the evoked spike times relative to each mechanical stimulation marker (−2 to +5 seconds around mechanical stimulation (Figure 2F)). Mechanically evoked spikes were extracted using the MATLAB function ‘histcounts’ to sum the total evoked spikes occurring in 200ms bins relative to each stimulation (−2 to +5 seconds peri-stimulus). A unit was classified as a wide dynamic range (WDR) neuron if it responded to dynamic brushing of the hind paw with maintained spike discharge, if it was activated at the onset and/or offset (transient) of punctate mechanical stimulation of the receptive field with von Frey filaments (with an increase in maintained spiking at von Frey filaments >15g), and if it produced maintained firing during a noxious pinch. Conversely, a unit was classified as a low threshold mechanoreceptive (LTMR) neuron if it fired at the onset and/or offset of the dynamic brush, and if it produced transient spiking to innocuous punctate mechanical stimulation of the RF with von Frey filaments (that did not increase at von Frey filaments >15g) or in response to noxious pinch.

The mechanically evoked spike counts for each unit was quantified by summing the number of spikes occurring from 0-3 seconds from the onset of the mechanical stimuli (brush, graded von Frey filaments, and pinch). This was averaged across repeated mechanical stimuli of each intensity (except pinch) within each period of the recording (baseline, tapentadol, and naloxone/ atipamezole), to generate an averaged evoked spike count for each unit.

The latency of the unit’s spike times evoked by electrical stimulation was used to estimate the likely classes of primary afferent fibre being activated by the electrical stimuli and mediating the spinal neuronal responses. Raster plots were used to visualise spiking latency following electrical stimulation of the sciatic nerve (0.1 second post-stimulus for 4Hz low-intensity stimulation and 0.3 seconds post-stimulus for Inn-Nox stimulation). The electrically evoked spikes were extracted using the MATLAB function ‘histcounts’ to sum the total evoked spikes occurring in 10ms bins relative to each electrical stimulation. Spikes occurring between 5-10ms post-stimulation of the sciatic nerve (∼10cm from the recording site) had an estimated conduction velocity of >10m/s and considered to be mediated by Aβ-fibres. Spike times between 20-50ms post-stimulation had an estimated conduction velocity of 2-5m/s, and were attributed to Aδ-fibre mediated activity, whereas and spikes with a latency >50ms post-stimulation were estimated to have a conduction velocity <2m/s, and were considered as being C-fibre mediated [15]. Single unit spike counts evoked by 4Hz electrical stimulation were calculated by averaging the number of evoked spikes across repetitive stimuli within each stimulus block and then averaging across blocks within each experimental period. Single unit spike counts evoked by the Inn-Nox electrical stimulation were averaged across repeated stimuli at each intensity within each experimental period. This generated averaged evoked spike counts of Aβ-fibre mediated activity to the 4Hz electrical stimulation, and averaged evoked spike counts of Aβ-, Aδ-, and C-fibre mediated activity evoked by the Inn-Nox electrical stimulation (at each stimulus intensity), at baseline, after tapentadol and following naloxone or atipamezole.

#### Analysis of somatosensory evoked potentials

To analyse the SEPs, all recording data and electrical stimulus time stamps were imported into MATLAB. Data were low pass filtered at 300Hz to remove stimulus artefacts. A peri-stimulus sample (−0.1 to +0.2s) was extracted from the data recorded from each electrode contact to isolate waveforms of the SEPs. These data samples were averaged across repeated stimuli within each stimulation block, and then across stimulation blocks in each experimental period, to produce a grand average SEP waveform for the baseline, tapentadol and naloxone / atipamezole periods for all channels. The amplitude of the N1 component recorded from each channel was extracted as the maximal primary negative peak of the SEP waveform between 0-10ms post stimulation. The peak amplitude value of the N1 component was taken from the channel with the largest signal.

Heatmap visualisations of the spinal SEPs recorded along the length of the probe were generated by first aligning the channel which recorded the maxima of the N1 component across animals. All 64-channels of data were then averaged across animals, to produce an averaged heatmap of the N1 component at baseline, in the presence of tapentadol, and following naloxone or atipamezole.

### Statistics

All statistical analysis of the effects of tapentadol, naloxone and atipamezole on single unit activity and SEP waveform amplitude were performed in GraphPad Prism (v10.6.1). Single unit evoked spike counts to electrical and mechanical stimulation are presented as PSTH, scatter plots and violin plots, as mean spike counts ±SEM. Statistical significance for the effect of tapentadol, naloxone and atipamezole on evoked activity in WDR and LTMR neurons (in response to electrical stimulation and mechanical stimulation) was calculated using a repeated measures one-way or two-way ANOVA depending on the data set, or using mixed effects analysis where data was missing due to the omission of a stimulus paradigm during an experiment. *Post-hoc* comparisons of single unit spike counts between functional classes and experimental periods was calculated using Tukey’s multiple comparisons test. A student’s t-test was used to assess differences in the depth profiles within the spinal cord of the functional populations.

SEP data are presented as mean waveforms and SEP component peak amplitude ±SEM. Statistical significance for the effect of tapentadol, naloxone and atipamezole on the SEP component amplitudes was calculated using a repeated measures one-way ANOVA, with Tukey’s multiple comparisons test being used for *post-hoc* analysis. A Pearson’s r calculation was performed to determine the correlation between the change in WDR neuronal activity (% of baseline, single units averaged within animal) and change in the amplitude of the N1 component (% of baseline) following injection of tapentadol.

## Results

### Identification of WDR and LTMR neurons

Clustering the data from the spinal multielectrode recordings (n=12 rats) yielded a total of 59 well-isolated single units that showed both mechanically-evoked activity from the hindpaw and electrically-evoked activity from sciatic stimulation across the entire duration of the experiment (∼2 hours). Functional classification of these cells identified two populations of spinal neurons, WDR neurons (n=47 cells, 80%) and LTMR neurons (n=12 cells, 20%), on the basis of distinctive response profiles to mechanical stimulation (Figure 3). All WDR neurons showed graded firing patterns to mechanical stimulation, exhibiting sustained firing to dynamic brush and transient (ON and OFF) firing to innocuous von Frey stimulation (2g), which increased with increasing von Frey filament force (8g and 26g). They also showed maintained firing in response to noxious pinch (Figure 3A). Conversely, LTMR neurons demonstrated transient dynamic responses to brush, and only displayed transient onset / offset firing in response to von Frey filament simulation, which did not adapt to increases in filament weight at the higher forces. They did not show a maintained response to noxious pinch, again only firing at the onset and offset of the pinch (Figure 3B). Analysis of the spatial distribution of WDR neurons (putatively spanning dorsal horn laminae III-VI) and LTMR neurons (spanning dorsal horn laminae II-V) that the WDR neurons tended to be located deeper in the dorsal horn (WDR neuron depth 463 ± 23 µm vs LTMR neuron depth 372 ± 43 µm; Unpaired two-tailed t-test, t = 1.766, DF = 57, P = 0.082; Figure 3C).

**Figure 3.**
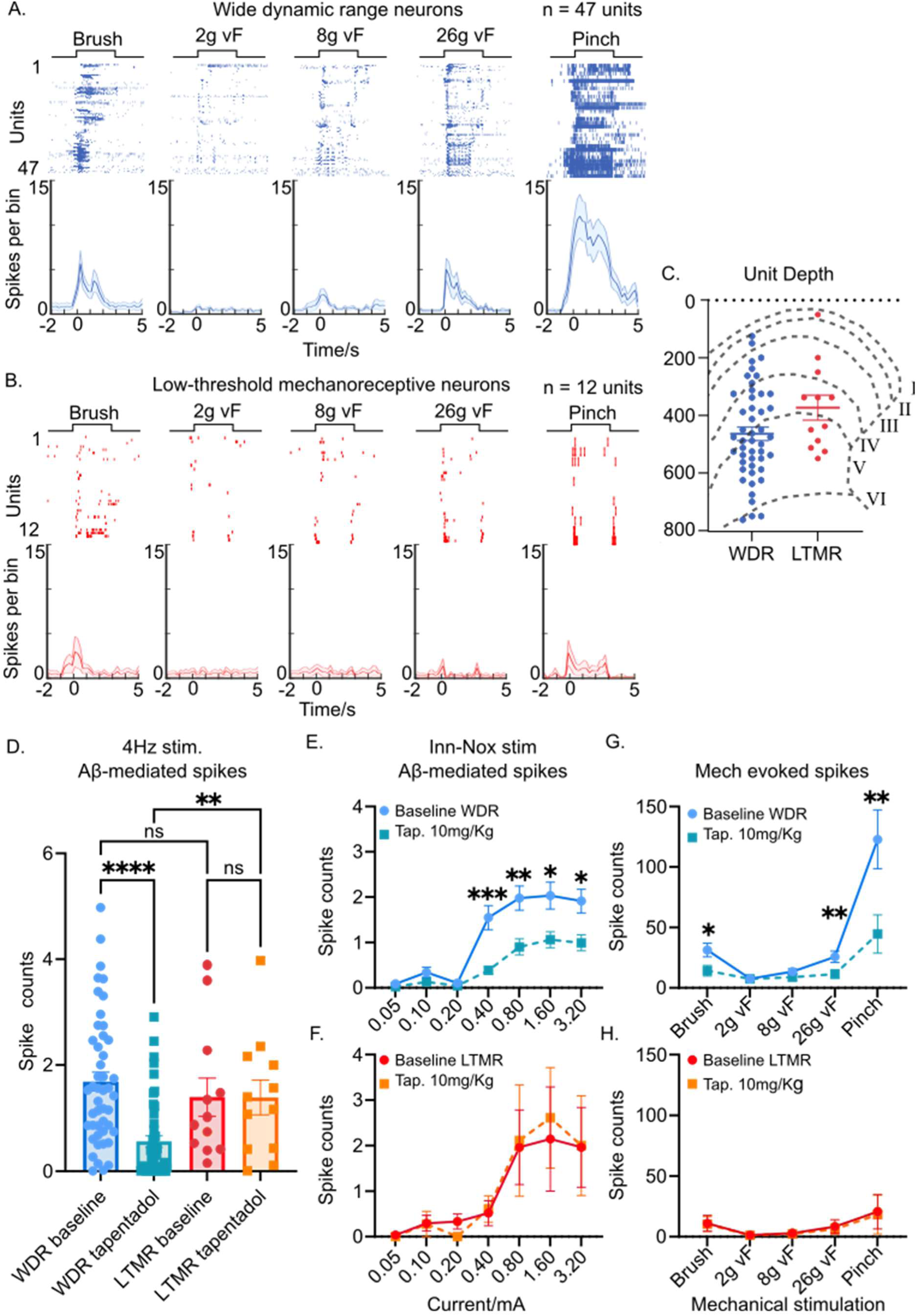
Selective effects of tapentadol on evoked activity of WDR and LTMR neurons. A. Spike raster plots and peristimulus time histograms of WDR neurons and B. LTMR neurons responses to mechanical stimulation of the hind paw. Single unit responses of 47 WDR neurons and 12 LTMR neurons were recorded from a cohort of 12 rats. C. The depth distribution of WDR neurons and LTMR neurons mapped to their approximate spinal dorsal horn laminae. D. Tapentadol selectively inhibits the Aβ-mediated activity of WDR neurons (compared to baseline responses to 4Hz electrical stimulation), without modulating LTMR neurons. Additionally, the Aβ-mediated activity, WDR neurons is significantly reduced compared to LTMR neurons, in the presence of tapentadol. E. Tapentadol significantly reduced the number of Aβ-mediated spikes during the Inn-Nox electrical stimulation (at intensities ≥0.4mA), F. Tapentadol had no effect on LTMR neuron Aβ-mediated activity. G. Tapentadol significantly reduced the spiking activity of WDR neurons in response to brush, 26g von Frey filament, and noxious pinch, H. Tapentadol did not change LTMR neurons responses to mechanical stimuli. Data presented as mean ±SEM.

### Tapentadol inhibits Aβ-fibre mediated activity in WDR but not LTMR neurons

As the N1 component of spinal SEPs is generated by the depolarisation of second order neurons and is attenuated by tapentadol [37], we investigated the effects of tapentadol on the Aβ-fibre mediated activity of WDR and LTMR neurons during the 4Hz (Low intensity) electrical stimulation protocol. At baseline, we saw no difference in the number of Aβ-latency evoked spikes between the WDR and LTMR neuronal populations (1.69 ± 0.18 vs 1.40 ± 0.35 spikes; mixed-effects analysis with Tukey’s multiple comparisons test: *q* = 1.323, DF = 11, *P* = 0.7868; Figure 3D). However, administration of tapentadol significantly reduced the Aβ-mediated activity of WDR neurons compared to baseline (1.69 ± 0.2 vs 0.56 ± 0.1 spikes, n=47; Tukey’s multiple comparison test: *q* = 9.186, DF = 46, *P* ≤ 0.0001; Figure 3D). Conversely, the Aβ-fibre latency evoked spike counts of LTMR neurons were not altered by tapentadol (1.40 ± 0.4 vs 1.39 ± 0.3 spikes, n=12; *q* = 0.03541, DF = 11, *P* > 0.9999). Accordingly, tapentadol significantly reduced Aβ-mediated WDR neuron activity compared to LTMR neurons (0.56 ± 0.1 vs 1.40 ± 0.4 spikes; *q* = 6.025, DF = 11, *P* = 0.0063).

We also analysed the effect of tapentadol on the Aβ-mediated activity of WDR and LTMR neurons during the Inn-Nox electrical stimulation, to determine how the drug modulates neuronal responses from the innocuous to the noxious range. This revealed a significant effect of tapentadol (baseline vs tapentadol; DF(3,84) = 4.213, P = 0.0079) and of stimulus intensity (DF(1.75,147.0) = 32.49, P < 0.0001) on the number of evoked spikes. Tapentadol selectively inhibited the electrically evoked activity of WDR neurons at stimulus intensities 0.4, 0.8, 1.6 and 3.2mA (Figure 3E), but not in response to lower intensities between 0.05-0.2mA (Tukey’s multiple comparisons test, Table 1). Conversely, tapentadol had no effect on the Aβ-mediated activity of LTMR neurons across all stimuli (Tukey’s multiple comparisons test, Figure 2F & Table 1). These findings demonstrate that tapentadol selectively inhibits the Aβ-mediated activity of WDR neurons during innocuous and noxious electrical stimulation, without modulating the activity of LTMR neurons.

**Table 1.** The effect of tapentadol on the Aβ-mediated activity of spinal neurons evoked by Inn-Nox electrical stimulation. Number of electrically evoked spikes of WDR and LTMR neurons (Mean ± SEM). Two-way ANOVA with Tukey’s multiple comparisons test.

| Stimulus (mA) | Baseline spikes (mean ± SEM) | Tapentadol spikes (mean ± SEM) | P value (Baseline vs Tapentadol) |
| --- | --- | --- | --- |
| **WDR neuronal responses to Inn-Nox stimulation (mA)** |  |  |  |
| 0.05 | 0.09±0.0 | 0.03±0.0 | 0.111 |
| 0.1 | 0.34 ±0.1 | 0.13 ± 0.0 | 0.310 |
| 0.2 | 0.1 ± 0.0 | 0.05 ± 0.0 | 0.540 |
| 0.4 | 1.55 ± 0.3 | 0.39 ± 0.1 | 0.011* |
| 0.8 | 1.98 ± 0.3 | 0.90 ± 0.2 | 0.007* |
| 1.6 | 2.03 ± 0.2 | 1.06 ± 0.2 | 0.035* |
| 3.2 | 1.91 ± 0.3 | 0.99 ± 0.2 | 0.024* |
| **LTMR neuronal responses to Inn-Nox stimulation (mA)** |  |  |  |
| 0.05 | 0.00 ± 0.1 | 0.00 ± 0.0 | 0.754 |
| 0.1 | 0.30 ± 0.2 | 0.19 ± 0.2 | 0.970 |
| 0.2 | 0.33 ± 0.2 | 0.00 ± 0.0 | 0.264 |
| 0.4 | 0.52 ± 0.3 | 0.41 ± 0.2 | 0.988 |
| 0.8 | 1.96 ± 0.8 | 1.41 ± 0.9 | 0.997 |
| 1.6 | 2.15 ± 1.1 | 1.74 ± 0.8 | 0.991 |
| 3.2 | 1.96 ± 0.9 | 1.33 ± 0.8 | 0.949 |

### Tapentadol produces a selective inhibition of WDR but not LTMR neuronal responses to intense mechanical stimuli

To test whether tapentadol differentially modulates mechanically evoked activity in WDR and LTMR neurons, we compared number of spikes evoked by brush, graded von Frey filaments, and pinch at baseline and 60 minutes after tapentadol administration. Mixed-effects analysis revealed there was a significant effect of tapentadol (compared to baseline; DF(1.392, 48.73) = 9.749, P = 0.0011) and stimulation modality (DF(1.087, 38.04) = 6.272, P = 0.0147) on the number of evoked spikes, and an interaction between these two factors (DF(1.621, 22.96) = 7.064, P = 0.0063), consistent with a differential modulation of WDR and LTMR neuron response profiles by tapentadol.

Post-hoc analysis showed that tapentadol attenuated WDR neuron responses in response to 26g von Frey filament and pinch (considered to be noxious), but not responses evoked by 2g or 8g von Frey filament (Figure 3G, Table 2). Tapentadol also significantly reduced the number of spikes evoked by brush, a finding of note given that tapentadol was not expected to modulate responses to innocuous stimuli, but consistent with its modulation WDR neuronal responses to 4Hz stimulation. In contrast, no significant tapentadol-induced change was observed in LTMR neuron activity for any stimulus modality (Figure 3H, Table 2). These findings demonstrate that tapentadol selectively attenuates the mechanically-evoked activity of WDR neurons, but not LTMR neurons, in response to both innocuous and noxious mechanical stimulation (as was the case with their electrically evoked activity).

**Table 2.** The effect of tapentadol on mechanically evoked activity of spinal neurons. Number of mechanically evoked spikes of WDR and LTMR neurons (Mean ± SEM). Two-way ANOVA with Tukey’s multiple comparison test.

| Stimulus | Baseline spikes (mean ± SEM) | Tapentadol spikes (mean ± SEM) | P value (Baseline vs Tapentadol) |
| --- | --- | --- | --- |
| **WDR neuronal responses to mechanical stimulation** |  |  |  |
| Brush | 31.30 ± 5.65 | 14.18 ± 4.34 | 0.009* |
| 2g von Frey | 7.53 ± 1.68 | 7.45 ± 2.99 | > 0.999 |
| 8g von Frey | 13.48 ± 2.33 | 8.88 ± 2.97 | 0.364 |
| 26g von Frey | 25.74 ± 4.64 | 11.36 ± 3.53 | 0.002* |
| Pinch | 122.78 ± 24.34 | 44.53 ± 15.82 | 0.001* |
| **LTMR neuronal responses to mechanical stimulation** |  |  |  |
| Brush | 11.00 ± 6.53 | 10.42 ± 6.09 | 0.999 |
| 2g von Frey | 1.36 ± 0.58 | 1.12 ± 0.74 | 0.963 |
| 8g von Frey | 2.91 ± 1.51 | 2.27 ± 1.01 | 0.888 |
| 26g von Frey | 8.33 ± 5.71 | 6.24 ± 4.17 | 0.687 |
| Pinch | 20.73 ± 14.22 | 18.27 ± 15.92 | 0.968 |

### Naloxone recovers the tapentadol-mediated inhibition of activity in WDR neurons

To investigate the receptor mechanisms underpinning the inhibitory effect of tapentadol on WDR neuronal activity and the amplitude of spinal SEPs, the animals were allocated into two cohorts which received either naloxone (0.25mg/Kg, i.p, n=5 rats) or atipamezole (1mg/Kg, i.p, n=7 rats), to attempt to reverse the µ-opioid receptor (MOR)-mediated and NRI-mediated effects of tapentadol, respectively. WDR neurons receive convergent synaptic inputs from multiple primary afferent fibres and are responsible for encoding the intensity and location of stimuli. Therefore, we assessed the effects of tapentadol and the antagonists on WDR neuronal activity mediated by Aβ-, Aδ- and C-fibre afferents based on their estimated response latencies and activation thresholds to electrical stimulation.

The Aβ-mediated activity of 22 WDR neurons (n=5 rats) was measured at baseline during the 4Hz stimulation, and again after subsequent injections of tapentadol (0-60 minutes), followed by naloxone (60-85 minutes). Tapentadol significantly inhibited the Aβ-fibre mediated firing of WDR neurons (one-way ANOVA with Tukey’s multiple comparisons test; 1.86 ± 0.3 vs 0.34 ± 0.1:*q* = 8.113, DF = 21, P<0.0001; Figure 4A) during the 4Hz stimulation paradigm. This inhibition was partially reversed by naloxone (0.34 ± 0.1 vs 1.06 ± 0.2 , *q* = 5.717, DF = 21, P = 0.0016), although this was still reduced compared to baseline level (one-way ANOVA with Tukey’s multiple comparisons test; 1.86 ± 0.3 vs 1.06 ± 0.2, *q* = 4.033, DF = 21, P = 0.0255; Figure 4A).

**Figure 4.**
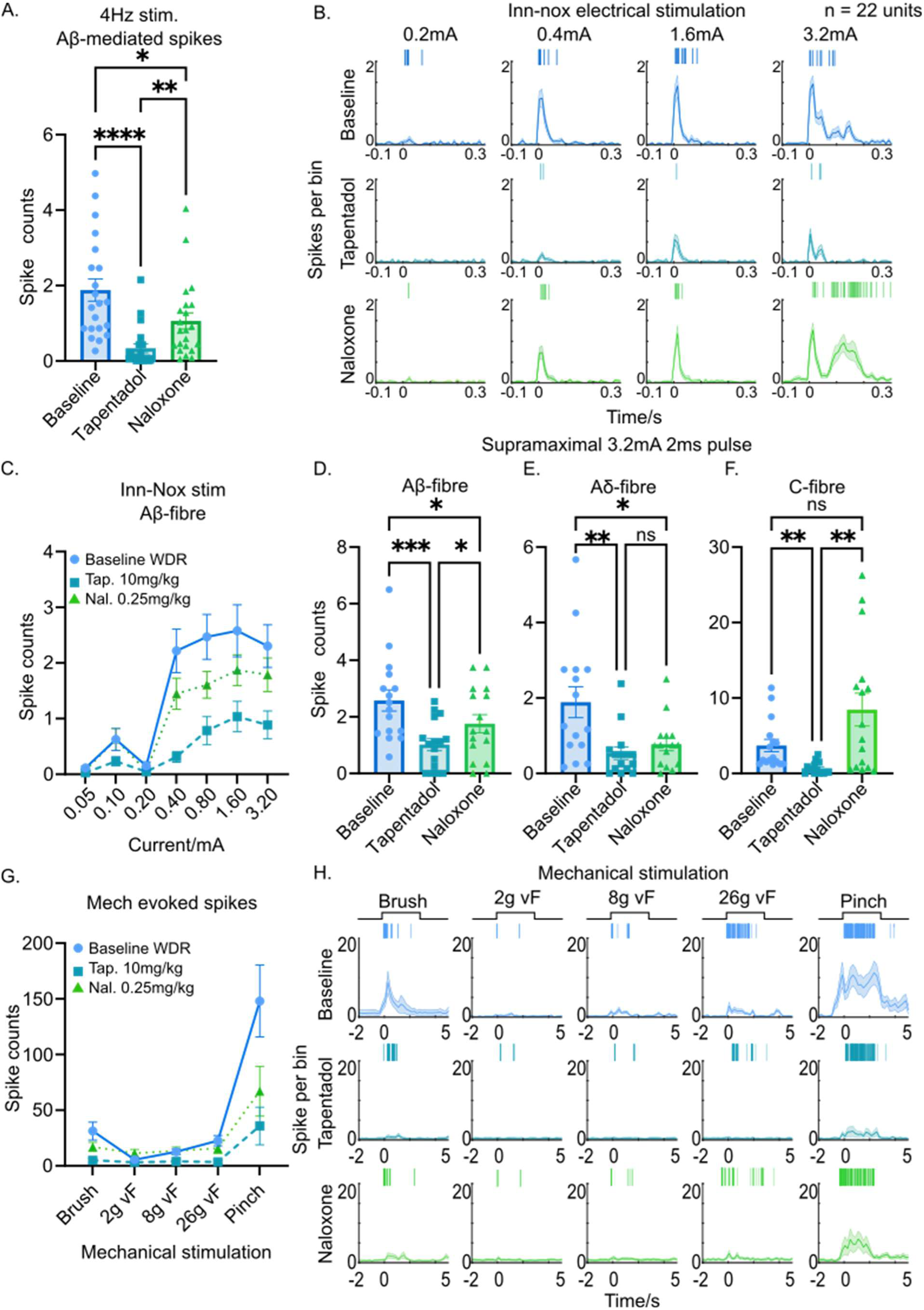
Naloxone recovers the inhibitory effects of tapentadol on WDR activity. A. The Aβ-mediated spike counts of WDR neurons in responses to innocuous 4Hz electrical stimulation. Naloxone partially reverses the inhibitory effects of tapentadol. B. PSTH of the electrically evoked activity of WDR neurons during the Inn-Nox stimulation at baseline (top, blue), in the presence of tapentadol (middle, green) and following injection of naloxone (bottom green). Accompanied with a representative raster plot of a representative single unit responses to each stimulus intensity. C. Stimulus response relationship showing Aβ-mediated spike counts of WDR neurons in response to Inn-Nox stimulation. Naloxone fully reversed the effects of tapentadol on WDR responses to 1.6mA and 3.2mA pulses, and partially reversed responses to 0.4 and 0.8mA pulses. D. The Aβ-, E. Aδ-, and F. C-fibre mediated activity of WDR neurons in responses to a supramaximal 3.2mA 2ms pulse. Naloxone partially reversed the effects of tapentadol on the Aβ-mediated activity of WDR neurons and fully reversed the inhibition of the C-fibre mediated activity but failed to recover Aδ-mediated responses of WDR neurons. G. The mechanically evoked spike counts of WDR neurons at baseline, in the presence of tapentadol and following injection of naloxone. Naloxone fully reversed the inhibitory effects of tapentadol on WDR responses to 8g and 26g von Frey filaments and pinch and partially recovered their responses to brush. H. PSTH of WDR neurons responses to mechanical stimulation during each experimental period, accompanied by a representative raster plot of a single units’ response to each stimulus. Data are presented as mean ±SEM. 22 WDR neurons were recorded from a cohort of 5 rats.

Tapentadol also significantly reduced the Aβ-mediated activity of WDR neurons evoked by the Inn-Nox stimulation protocol, at intensities ≥0.4mA (Figure 4B-C & Table 3). Naloxone fully reversed the evoked firing of WDR neurons in response to stimulation intensities of 1.6mA and 3.2mA and partially recovered WDR responses to 0.4 and 0.8mA stimulation (Table 3).

**Table 3.**
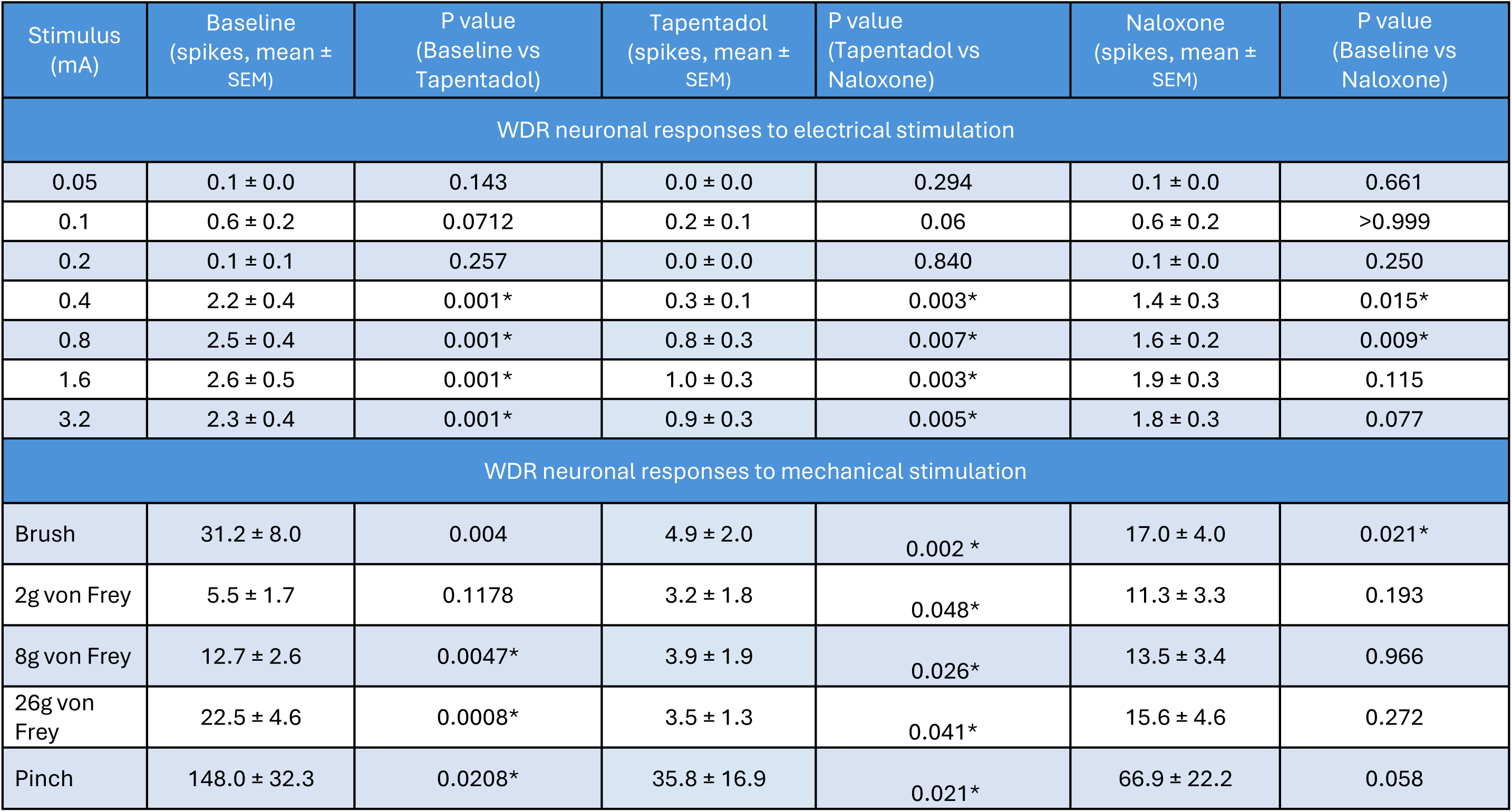
The effect of tapentadol and naloxone on WDR neuronal activity evoked by electrical and mechanical stimulation. Number of electrically and mechanically evoked spikes of WDR neurons (Mean ± SEM). Two-way ANOVA with Tukey’s multiple comparisons test.

To assess the effects of tapentadol and naloxone on the Aδ-fibre and C-fibre mediated activity of WDR neurons, we assessed the number of spikes evoked by a supramaximal electrical stimulus (3.2mA x 2ms to maximally recruit peripheral nociceptors), at latencies equivalent to the estimated conduction velocities of primary afferent fibre classes (Figure 4D-F). Compared to baseline responses to the supramaximal stimulation, tapentadol significantly reduced the number of Aβ-fibre (2.58 ± 0.4 vs 1.02 ± 0.2 spikes; *q* = 7.782, DF = 17, P = 0.0001), Aδ-fibre (1.89 ± 0.4 vs 0.53 ± 0.2 spikes; *q* = 5.220, DF = 17, P = 0.0064), and C-fibre (3.69 ± 0.8 vs 0.70 ± 0.2; repeated measures one-way ANOVA with Tukey’s multiple comparisons test; *q* = 5.47, DF = 17, P = 0.0041) mediated spikes (Figure 4D-F). Naloxone fully reversed the tapentadol-induced suppression of C-fibre mediated activity of WDR neurons (0.70 ± 0.2 vs 8.46 ± 2.2 spikes; *q* = 5.170, DF = 17, P = 0.0062) and partially reversed the effects of tapentadol on the Aβ-fibre mediated activity (1.02 ± 0.2 vs 1.76 ± 0.3 spikes; *q* = 3.796, DF = 17, P = 0.0424), however this was still significantly lower than baseline (1.76 ± 0.3 vs 2.58 ± 0.4 spikes; repeated measures one-way ANOVA with Tukey’s multiple comparisons test; *q* = 3.767, DF = 17, P = 0.0441) (Figure 4D-F). Naloxone did not significantly reverse the inhibitory effects of tapentadol on the Aδ-fibre mediated activity (0.53 ± 0.2 vs 0.77 ± 0.2;*q* = 2.104, P = 0.3262).

Tapentadol significantly reduced the number of spikes evoked in WDR neurons by brush, 8g and 26g von Frey filament, and pinch, but not to 2g von Frey filament (Mixed-effects model with Tukey’s multiple comparisons test; stimulus intensity DF(1.072, 21.44) = 13.19, P=0.0013; baseline vs tapentadol vs naloxone DF(1.350, 27.0) = 17.93, P < 0.0001; stimulus intensity x experimental period interaction DF(1.265, 22.45) = 7.205, P = 0.0093; Figure 4G-H & Table 3). Naloxone fully reversed the tapentadol-mediated inhibition of mechanically-evoked activity in response to 8g and 26g von Frey filaments, and to pinch back to baseline levels, and partially reversed WDR neurons responses evoked by brush (Figure 4G-H & Table 3).

These data demonstrate that naloxone recovers most of the inhibitory effects of tapentadol on WDR neuronal activity, mediated by Aβ-fibre and C-fibre primary afferents, and in a manner which reflects their encoding of high intensity mechanical stimulation.

### Atipamezole does not reverse the tapentadol-mediated inhibition of WDR neuronal responses

In addition to MOR agonism, tapentadol is also a noradrenaline reuptake inhibitor (NRI) acting on the transporter to increase the extracellular concentration of noradrenaline. We sought to investigate how this NRI activity contributes to the inhibition of WDR neuronal activity by using atipamezole to block the inhibitory α_2_-adrenoreceptor, which is thought to mediate the noradrenergic-inhibition of spinal neurons [43].

We analysed the behaviour of 25 WDR neurons (n=7 rats), evoked by 4Hz, Inn-Nox electrical stimulation and mechanical stimulation protocols, at baseline, and following injections of tapentadol followed by atipamezole (1mg/kg). Tapentadol again significantly reduced the Aβ-fibre mediated spike counts of WDR neurons during 4Hz electrical stimulation (1.54 ± 0.2 vs 0.75 ± 0.2 spikes; Repeated measures one-way ANOVA with Tukey’s multiple comparisons test; *q* = 5.279, DF = 24, P = 0.0029) when compared to baseline (Figure 5A). However, atipamezole did not significantly reverse the tapentadol inhibition of Aβ-fire mediated spiking of WDR neurons (0.75 ± 0.2 vs 1.06 ± 0.2 spikes; Tukey’s multiple comparisons test; *q* = 2.212, DF= 24, P = 0.2803) which was still significantly reduced compared to baseline (1.06 ± 0.2 vs 1.54 ± 0.2 spikes; Tukey’s multiple comparisons test; *q* = 4.052, DF = 24, P = 0.0224; Figure 5A).

**Figure 5.**
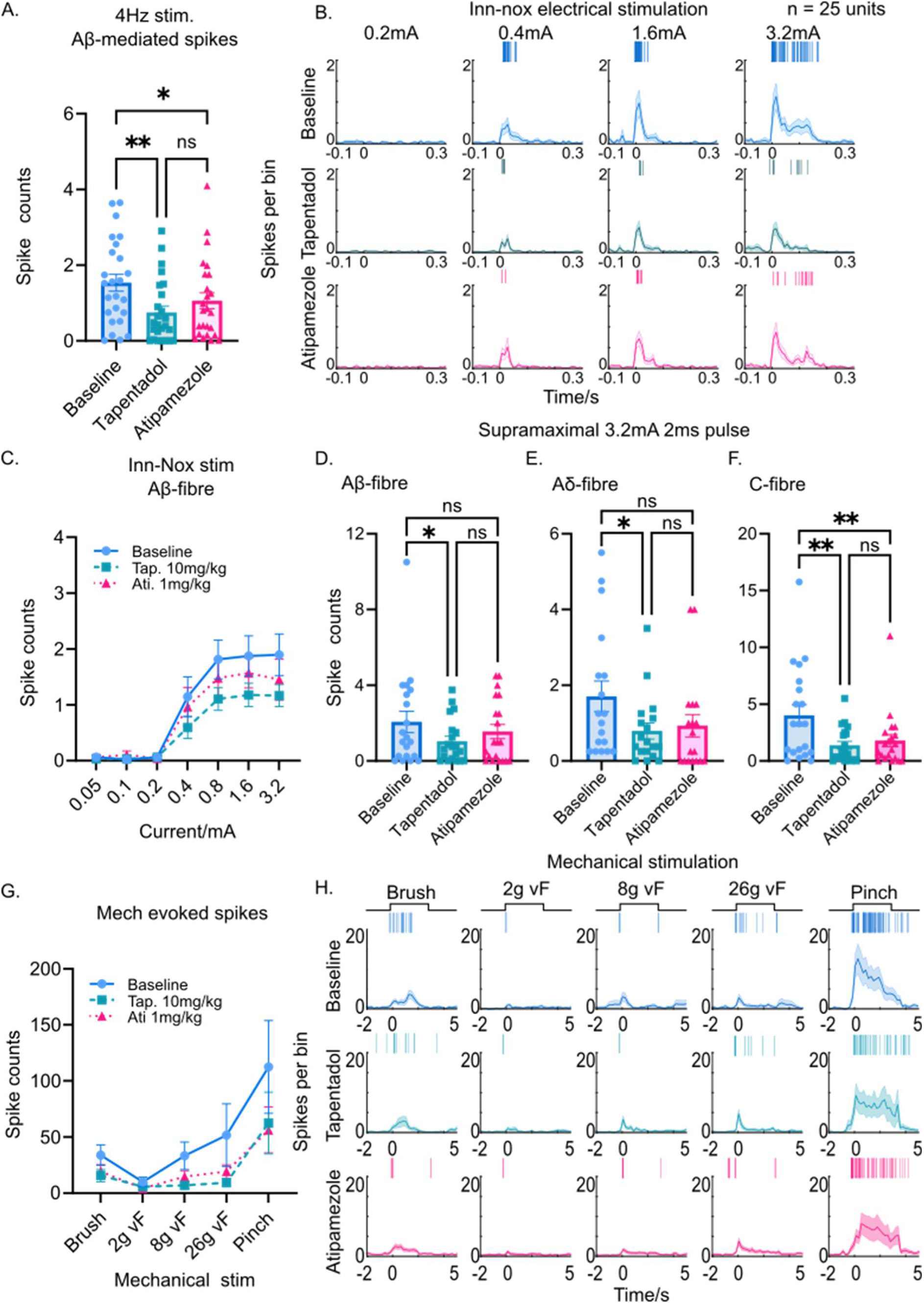
Atipamezole does not attenuate the inhibitory effects of tapentadol on WDR neuronal responses to electrical and mechanical stimulation. A.The Aβ-mediated spike counts of WDR neurons in responses to innocuous 4Hz electrical stimulation. Atipamezole did not reverse the inhibitory effects of tapentadol. B. PSTH of the electrically evoked activity of WDR neurons during the Inn-Nox stimulation at baseline (top, blue), in the presence of tapentadol (middle, green) and following injection of atipamezole (bottom magenta). Accompanied with a representative raster plot of a single units’ responses to each stimulus intensity. C. The Aβ-mediated spike counts of WDR neurons in response to Inn-Nox stimulation. Atipamezole failed to reverse the effects of tapentadol on WDR responses. D. The Aβ-, E. Aδ-, and F. C-fibre mediated activity of WDR neurons in responses to a supramaximal 3.2mA 2ms pulse. Atipamezole did not reverse the effects of tapentadol on the Aβ-, Aδ- and C-fibre mediated activity of WDR neurons. G. The mechanically evoked spike counts of WDR neurons at baseline, in the presence of tapentadol and following injection of atipamezole. Atipamezole did not reverse the inhibitory effects of tapentadol on WDR responses to any stimulus modality. H. PSTH of WDR neurons responses to mechanical stimulation during each experimental period, accompanied by a representative raster plot of a single units’ response to each stimuli. Data are presented as mean ±SEM.

During the Inn-Nox protocol, tapentadol significantly reduced the Aβ-fibre mediated spike counts of WDR neurons stimulus intensities of ≥0.4mA (Figure 5B-C & Table 4). However, atipamezole did not significantly reverse the inhibitory effects of tapentadol on the Aβ-fibre mediated spiking of WDR neurons at any stimulus intensity (Figure 5B-C & Table 4).

**Table 4.** The effect of tapentadol and atipamezole on WDR neuronal activity evoked by electrical and mechanical stimulation. Mean number of electrically and mechanically evoked spikes ± SEM of WDR neurons. Two-way ANOVA with Tukey’s multiple comparisons test.

| Stimulus (mA) | Baseline (spikes, mean ± SEM) | P value (Baseline vs Tapentadol) | Tapentadol (spikes, mean ± SEM) | P value (Tapentadol vs Atipamezole) | Naloxone (spikes, mean ± SEM) | P value (Baseline vs Atipamezole) |
| --- | --- | --- | --- | --- | --- | --- |
| WDR neurons' responses to electrical stimulation |  |  |  |  |  |  |
| 0.05 | 0.1 ± 0.0 | 0.98 | 0.0 ± 0.0 | 0.987 | 0.1 ± 0.0 | >0.999 |
| 0.1 | 0.0 ± 0.0 | 0.999 | 0.0 ± 0.0 | 0.93 | 0.1 ± 0.1 | 0.910 |
| 0.2 | 0.1 ± 0.0 | >0.999 | 0.1 ± 0.0 | 0.994 | 0.0 ± 0.0 | 0.909 |
| 0.4 | 1.1 ± 0.4 | 0.019* | 0.6 ± 0.2 | 0.158 | 1.0 ± 0.3 | 0.642 |
| 0.8 | 1.8 ± 0.3 | 0.002* | 1.1 ± 0.2 | 0.153 | 1.5 ± 0.3 | 0.225 |
| 1.6 | 1.9 ± 0.4 | 0.002* | 1.2 ± 0.2 | 0.128 | 1.6 ± 0.3 | 0.286 |
| 3.2 | 1.9 ± 0.4 | 0.001* | 1.2 ± 0.2 | 0.335 | 1.5 ± 0.4 | 0.073 |
| WDR neurons' responses to mechanical stimulation |  |  |  |  |  |  |
| Brush | 33.8 ± 9.0 | 0.175 | 15.6 ± 5.5 | 0.935 | 19.2 ± 6.1 | 0.935 |
| 2g von Frey | 10.0 ± 4.4 | 0.909 | 5.7 ± 3.1 | 0.986 | 4.1 ± 1.7 | 0.832 |
| 8g von Frey | 33.3 ± 12.1 | 0.028* | 6.9 ± 3.2 | 0.731 | 14.6 ± 5.4 | 0.159 |
| 26g von Frey | 51.5 ± 27.9 | 0.0002* | 9.2 ± 3.6 | 0.591 | 19.2 ± 6.0 | 0.005* |
| Pinch | 112.3 ± 41.4 | <0.0001* | 62.4 ± 27.5 | 0.813 | 56.2 ± 20.4 | <0.0001* |

Next, we investigated whether atipamezole reversed any fibre-specific modulatory effects of tapentadol on WDR neuronal activity, by analysing the number of spikes evoked by a supramaximal 3.2mA x 2ms pulse, at Aβ-, Aδ- and C-fibre equivalent latencies. Tapentadol again significantly reduced the number of Aβ-fibre mediated (2.06 ± 0.6 vs 1.04 ± 0.3 spikes; *q* = 3.717, DF = 19, P = 0.0419), Aδ-fibre mediated (1.71 ± 0.4 vs 0.79 ± 0.2 spikes; *q* = 4.329, DF = 19, P = 0.0184), and C-fibre mediated spikes (4.03 ± 0.9 vs 1.38 ± 0.3 spikes; Repeated measures one-way ANOVA; *q* = 5.124, DF = 19, P = 0.0049). However, injection of atipamezole, did not significantly reverse this inhibition of Aβ-fibre (1.04 ± 0.3 vs 1.55 ± 0.4;*q* = 2.551, DF = 19, P = 0.1951), Aδ-fibre (0.79 ± 0.2 vs 0.93 ± 0.3 spikes; *q* = 0.7, DF = 19, P = 0.8746) or C-fibre (1.38 ± 0.3 vs 1.79 ± 0.6 spikes; *q* = 1.315, DF = 19, P =0.6285) mediated latencies and the C-fibre mediated spiking of WDR neurons remained significantly reduced compared to baseline (1.79 ± 0.6 vs 4.03 ± 0.9; *q* = 2.859, DF = 19, P = 0.0074) (Figure 5D,E,F).

Finally, we also analysed whether atipamezole reversed the inhibitory effects of tapentadol on the WDR neuron’s responses to mechanical stimulation. Tapentadol again significantly reduced the number of spikes evoked by 8g and 26g von Frey filaments and by pinch, but not in response to brush or 2g von Frey filaments (Figure 5). Atipamezole did not significantly reverse the inhibitory effects of tapentadol on WDR neuronal responses to 8g and 26g von Frey filaments or pinch, which were still significantly reduced when compared to baseline.

In contrast to naloxone, these data show that atipamezole does not reverse the inhibitory effects of tapentadol on WDR neuronal activity evoked by mechanical or electrical stimulation. Together, these findings suggests that tapentadol exerts its inhibitory effects on WDR neurons predominantly via a MOR-mediated mechanism.

### The amplitude of the N1 SEP correlates with the inhibition and recovery of WDR neuronal activity during 4Hz electrical stimulation

The selective inhibition of WDR neurons, but not LTMR neurons, by tapentadol supported the hypothesis that the N1 wave might contain a component that reflected WDR activity and therefore serve as a translatable metric of nociceptive circuit excitability. To further examine this hypothesis, we investigated the relationship between the inhibitory effects of tapentadol on WDR neuronal excitability and N1 amplitude. We analysed the amplitude of the N1 component of the spinal SEPs generated during the 4Hz electrical stimulation at baseline and following i.p injections of tapentadol and naloxone/atipamezole (Figure 6).

**Figure 6.**
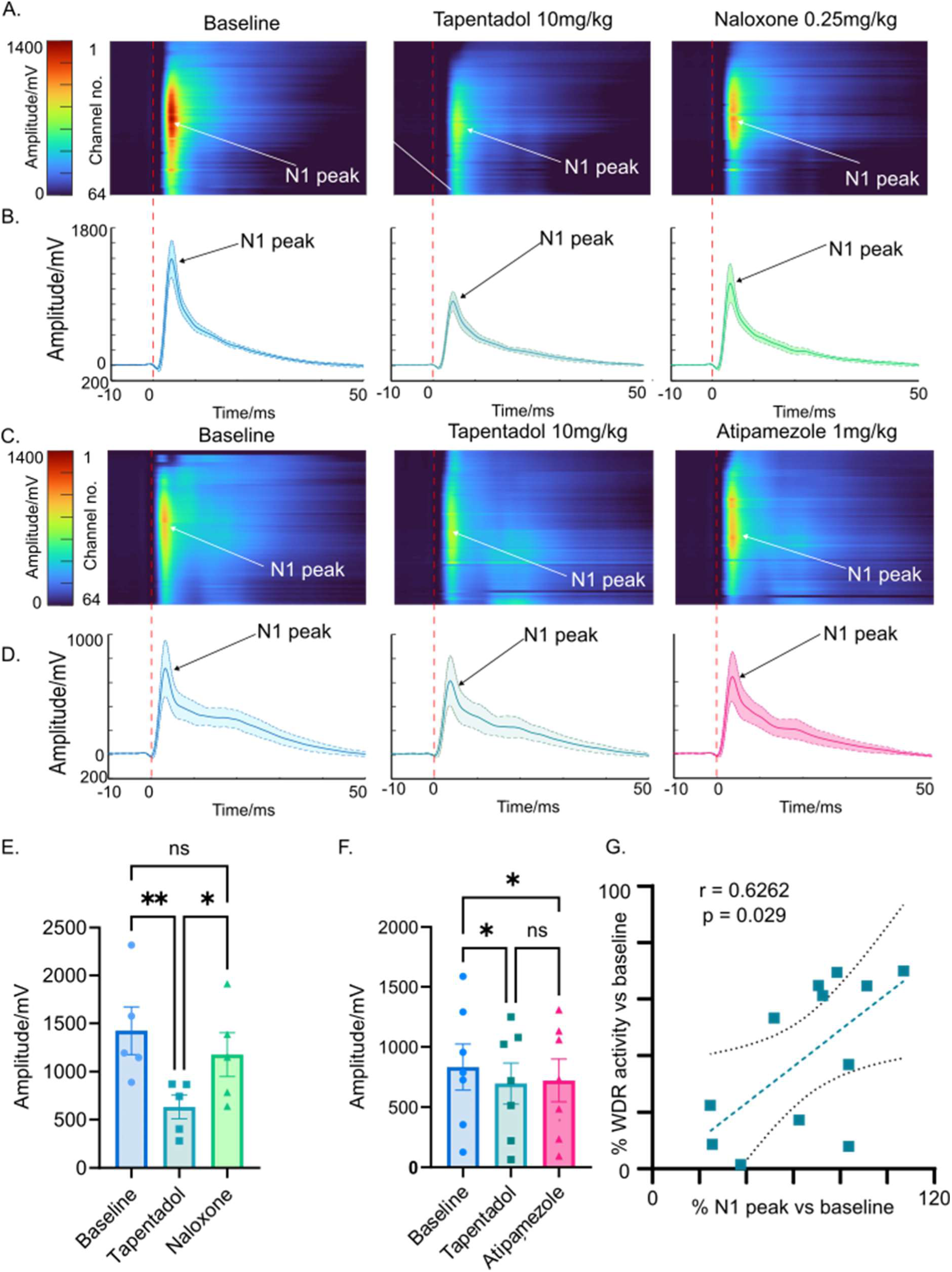
Amplitude of the N1 SEP correlates with the activity of WDR neurons and their modulation by tapentadol, naloxone and atipamezole. A.Heatmap visualisations of spinal SEP generated during the 4Hz electrical stimulation paradigm. Heatmaps show the averaged spinal SEP from 5 rats, recorded at different depths from each channel of the multielectrode probe (1-64) as it evolves over time, at baseline, in the presence of tapentadol and following injection of naloxone. Arrows point to the peak of the N1 component, B. which was extracted from the multielectrode data to analyse changes in the amplitude of the evoked potential. C. Heatmap visualisations show the averaged spinal SEP recorded from 7 rats across the multielectrode probe at baseline, in the presence of tapentadol and following injection of atipamezole. D. The averaged trace capturing the peak amplitude of the N1 component during each experimental period. E. Tapentadol significantly reduced the peak amplitude of the N1 component compared to baseline, and this was reversed by naloxone. F. Atipamezole failed to reverse the inhibitory effects of tapentadol on the amplitude of the N1 component. G. Scatterplot of the change in the averaged spike counts of WDR neurons vs the change in amplitude of the N1 SEP (within each animal, n = 12), generated during the 4Hz electrical stimulation, as a % of baseline. A linear regression generated a line of best fit with 95% confidence intervals and the Pearson’s correlation coefficient r = 0.6276, P = 0.029.

Spinal SEPs were recorded across all channels of the multielectrode probe. Heatmap visualisations show that the primary N1 component occurred ∼4ms post-stimulation, suggesting a conduction velocity of >25m/s (Aβ-fibre), and was largest in the mid channels of the electrode (around dorsal horn laminae IV-V; Figure 6A-D). Tapentadol significantly reduced the peak amplitude of the N1 evoked during the 4Hz stimulation paradigm (one-way ANOVA with Tukey’s multiple comparisons test; 1424 ± 248 µV vs 632 ± 122 µV.; *q* = 5.897, DF = 8, P = 0.0078), and this effect was reversed by naloxone (632 ± 122 µV vs 1175 ± 226 µV; *q* = 4.044, DF = 8, P = 0.0498. Figure 6E), just as naloxone reversed the inhibition of WDR neuronal activity by tapentadol.

In the rats which received injections of tapentadol and atipamezole, again the amplitude of the N1 component evoked during the 4Hz stimulation was shown to be significantly reduced by tapentadol (one-way ANOVA with Tukey’s multiple comparisons test; 832 ± 191 µV vs 695 ± 170 µV; *q* = 5.897, DF = 12 P = 0.0078, Figure 6F). However, atipamezole did not significantly recover the amplitude of the N1 SEP during the 4Hz stimulation (695 ± 170 µV vs 721 ± 176 µV; *q* = 0.8926, DF = 12, P = 0.8926, Figure 6F).

Across all animals (n = 12), the % change in amplitude of the N1 component recorded during the tapentadol period (/baseline *100) showed a positive correlation with the average % change in spike counts of all WDR neurons within each animal evoked during the 4Hz electrical stimulation (Pearson’s correlation coefficient; r = 0.6276, P = 0.0289, Figure 6G). This indicated there was a moderate to strong relationship between the changes in SEP amplitude and WDR neuronal excitability during the 4Hz electrical stimulation.

## Discussion

The innocuously-evoked N13 component of human spinal SEPs has been proposed as a translatable biomarker of spinal nociceptive processing, capable of demonstrating analgesic target engagement at the spinal level. The neuronal correlates of this evoked potential had never been defined, and it was unclear whether they were involved in the encoding of nociceptive stimuli and hence sensitive to modulation by analgesics. Here, we have shown that the activation of both WDR and LTMR neurons contributes to the generation of the N1 component. However, it is the selective inhibition of WDR neurons, but not LTMR neurons, by tapentadol, which attenuates the amplitude of the N1 component in naïve rats. This effect is predominantly mediated via a MOR-mediated mechanism of action and suppresses the evoked activity of WDR neurons to stimuli across the innocuous-to-noxious range. Moreover, the degree of inhibition of WDR neurons by tapentadol correlates with the change in amplitude of the N1 component. Taken together, this demonstrates that a substantial component the N1 potential is reflective of changes in sensory processing by WDR neurons, to both innocuous and noxious stimuli.

The use of multielectrode recordings with *post-hoc* spike sorting has emerged as a powerful technique for electrophysiological recordings, combining the specificity of single-unit recordings that enable the interrogation of single cell behaviours with the breadth of field potential recordings which capture mass network changes in excitability [18]. We classified 59 simultaneously recorded spinal dorsal horn neurons from 12 animals into functionally distinct populations of WDR neurons (predominantly in lamina IV-V) or LTMR neurons (predominantly in lamina III-IV). The spatial distribution of our LTMR neurons and WDR neurons is consistent with previous studies, respectively, defining the functional organisation of LTMR neurons in laminae III-V [24], and capturing the encoding of mechanical stimuli from lamina V WDR neurons [40].

WDR neurons receive convergent inputs from multiple primary afferent fibres, to encode the intensity of a stimulus, some of these neurons then transmit this information to the brain via ascending axonal projections [40]. LTMR neurons receive monosynaptic inputs from type I and type II rapidly adapting Aβ-fibres and are responsible for encoding the onset / offset of light touch [2,24]. We provide spatial and temporal evidence demonstrating that WDR and LTMR neurons are both synchronously activated by the innocuous electrical stimulation, during the generation of the N1 potential, and that their distribution within the dorsal horn mirrors the location of the spinal evoked potential. This first volley of evoked activity occurs <10ms after sciatic nerve stimulation consistent with Aβ-fibre mediated transmission with conduction velocities of >20m/s [1,24]. These findings provide evidence that the Aβ-mediated activation of WDR neurons, along with LTMR neurons, is central to the generation of the N1 component of spinal SEPs.

The inhibition of WDR neurons by tapentadol is likely mediated by both pre-synaptic inhibition of afferent nociceptors, which express the MORs [7,21], and post-synaptic inhibition of WDR neurons, as the responses of WDR neurons to Aβ-fibres (which do not express MORs) were also inhibited. The inhibition of the Aβ-mediated activity of WDR neurons by tapentadol showed a moderate-to-strong correlation with amplitude of the N1 potential. This finding addresses our first question, by demonstrating that the inhibition of the neuronal correlates of spinal SEPs leads to an attenuation of the amplitude of the N1 component. We have also shown that the responses of WDR neurons to noxious stimuli are inhibited by tapentadol and selectively recovered by naloxone, indicating MOR activation as the predominant effector. This second finding is evidence of analgesic target engagement with and the inhibition of nociceptive processing by WDR neurons. In contrast, the responsiveness of LTMR neurons to all stimulus modalities was unaffected by tapentadol, an expected finding given that there is no clinical evidence of tapentadol causing numbness in patients [20]. The maintained activity of LTMR neurons also indicates that the inhibition of WDR neurons by tapentadol was not reflective of a total decrease in network neuronal excitability, or indeed a deterioration in spinal cord viability caused by the drug. Collectively, this evidence provides proof-of-principle that the innocuously-evoked N1 potential can be used to assay changes of nociceptive processing by WDR neurons in the spinal dorsal horn – as was previously postulated [22,23,25,30].

Whilst we are confident that the inhibition of WDR neurons by tapentadol was predominantly mediated by activation of the MOR, it was surprising that atipamezole failed to recover the activity of WDR neurons or the amplitude of the N1. Previous spinal electrophysiological recordings have demonstrated that the inhibition of WDR neurons by tapentadol can be reversed by naloxone and atipamezole in both sham and nerve injured rats [6,10]. It is however noteworthy that Bee et al [6] reported that atipamezole more efficaciously reversed the inhibition of mechanically evoked activity by tapentadol in nerve injured rats, whereas naloxone more potently reversed spinal neuronal responses to mechanical stimulation in naïve rats. This points to a shift towards noradrenergic mediated inhibition of nociceptive processing by tapentadol in neuropathic pain states. This is unlikely to be caused by local changes in MOR expression; a study of MOR immunoreactivity in the spinal dorsal horn reported no changes between naive and nerve injured rats [31]. We have previously shown that enhancing the actions of spinal noradrenaline is anti-hyperalgesic in behavioural experiments in neuropathic animals [17,19,36], and argued that the relative failure of descending noradrenergic control that can be mitigated by NRIs. It is the dual-action of tapentadol as a MOR-agonist and NRI that has seen it adopted as a treatment for neuropathic pain. [16,18,34]. However, the specific neuronal correlates and molecular mechanisms underpinning this alteration in descending noradrenergic control remain poorly understood – it is possible that spinal SEPs might serve as a useful tool to investigate the mechanisms of spinal noradrenergic analgesia and translate the findings between preclinical and clinical studies.

The use of spinal SEPs is currently receiving renewed interest, following the development of novel recording approaches which could potentially expand the capabilities of spinal SEPs as a clinical investigative tool [13,28]. The introduction of a surface multichannel electrode arrays positioned over the neck and trunk allows for the recording of the spatial distribution of cervical and lumbar SEPs, revealing between-participant differences in their spinal segmental generators [28]. Additionally, the incorporation of canonical correlation analysis into post-processing pipelines enabled the tailoring of analysis parameters to individual data sets, to maximise signal-to-noise ratios and SEP extraction. This technique allowed for the detection of changes in spinal SEPs evoked by the stimulation of different fingers, and the recording of ultra-late components of spinal SEPs, evoked nociceptive heat-pain induced by CO_2_ laser stimulation [28]. Whilst a multichannel recording approach enables a more precise study of the spatiotemporal dynamics of somatosensory in the human spinal cord, such an approach must now be validated and standardised if it is to replace single surface electrode recordings. Additionally, future clinical investigations of spinal SEPs must be guided by (or back translated post-hoc) into preclinical equivalent studies, capable of defining the neuronal circuit basis of spinal SEPs.

Whilst these experiments were performed in naïve rats, this assay should be repeated in pain models to determine whether differences in pain state affects the neuronal and field potential responses to the drugs. Additionally, there would be benefits to reproduce these findings in cohorts of different sexes and ages, as women are more likely to develop chronic pain conditions than men [3], and such conditions are more likely to occur in later life [11].

The spinal dorsal horn is a complex and heterogenous network of local interneurons and projection neurons and the WDR functional classification encompasses a wide variety of different cell classes [39]. Recent advances in single-cell RNA sequencing combined with spatial transcriptomics has redefined our understanding of the complexity of the spinal dorsal horn architecture [14,33]. Recent studies combining multielectrode recordings with genetically modified mouse lines have been able to attribute functional response characteristics to specific genetic markers of interneuron subtypes [8,32,41]. Future spinal electrophysiological studies will be needed to characterise the changes in responses of these genetically identified populations of interneurons and projection neurons to the array of sensory stimuli in both naïve and sensitised animal models, to better comprehend their roles in chronic pain disorders.

We have established that WDR neurons, in addition to LTMR neurons, contribute to the generation of the N1 component in response to innocuous electrical stimulation. Moreover, we have demonstrated that the inhibition of the N1 SEP by tapentadol correlates with the magnitude of inhibition of WDR neurons, identifying WDR neurons as the likely target of tapentadol, which act predominantly through MOR-activation in naïve rats. These findings provide proof-of-principle for how the N1 component of spinal SEPs could be used as reliable biomarker of spinal nociceptive processing, and a translatable metric of WDR neuronal excitability.

## Author Contributions

Kenneth Steel (Conceptualisation, Methodology, Validation, Formal analysis, Investigation, Data Curation, Writing – Original Draft, Writing – Review & Editing), Tony Blockeel (Conceptualisation, Methodology, Writing-Review & Editing, Supervision), Elise Ajay (Formal analysis, investigation, Data Curation, Writing – Review & Editing), Jeff Krajewski (Supervision, Writing – Review & Editing), Keith Phillips (Conceptualisation, Writing – Review & Editing, Supervision), James P Dunham (Investigation, Writing – Review & Editing, Supervision) and Anthony E Pickering (Conceptualisation, Resources, Writing – Review & Editing, Supervision, Funding Acquisition).

## Data availability

All data will be made available upon reasonable request.

